# Na/K pump: Single-channel vs. Two-access-channel

**DOI:** 10.1101/275263

**Authors:** P. Liang, J. Mast, W. Chen

## Abstract

We re-studied the dialyzed Na/K pumps, the experimental foundation for the two-access-channel model. We found that the charge-movement pump currents, the major evidence for the two-access-channel, are only observed in certain situations. Once the stimulation pulse is high enough at low ionic concentration gradient, the backward pump currents disappeared. The two-directional charge movement pump currents become uni-directional pump currents showing that ions have passed through the channel across the cell membrane.

A modified single-channel configuration is introduced to explain the pump currents as two-way charge movement current and the one-way current. The negatively charged amino acids deeply inside the pump channel constitute a spatial array that function as a collimator to align the moving ions in the channel lumen and an energy-well for the moving cations. If the stimulation pulse cannot drive ions to overcome the ionic concentration gradient, the ions will be entrapped into the energy-well as if the pump channel is obstructed showing two separated access-channels. Once the stimulation pulse is high enough to drive ions overcoming the ionic concentration gradient, ions will pass through the channel across the cell membrane so that Na/K pumps exhibit a single channel configuration.

## Introduction

Functions and structures of Na/K pump have been well studied. However, the mechanisms involved in the ion-transport across cell membrane remain unclear. Since Lauger first proposed the channel-like functions in the 1970’s (1), a significant effort has been committed to investigate the details of pump channel (2,3,4,5,6).

Studies of the Polytoxin-treated Na/K pumps showed a pathway from the extra-to intracellular solution (7,8,9). Cysteine-scanning mutagenesis studies of α-helices in transmembrane domain with MTSET or MTSES demonstrated that the amino acids in α-helices affecting the channel conductance are mainly located at the ends of channel (10-12). These studies indicated that Na/K pumps have a single channel configuration with orifices located at two ends of the channel. Synchronization of Na/K pumps (13) and modulation of the pumping rate (14) indicated that energy, instead of protein conformational changes, dominates the ion-movements through the channel which implies the single-channel configuration. Gadsby has postulated a model of the isolated single-channel configuration where Na and K ions freely move along the channel without the effects of ionic concentration gradient and the membrane potential difference (15).

On the other hand, studies of the partially dialyzed pump molecules have showed charge-movement-like pump currents that indicate the pump channel is obstructed deeply inside the membrane which separates the pump channel into two segments (access-channels) (16,17,18,19). Both the studies of polytoxin-treated pumps (7,8,9) and the molecular dynamics simulations (19) showed that the external access channel has a wide open vestibule. Na and K ions are bond and occluded at end of the corresponding access-channel, transported to another access-channel through the protein conformational changes, and then released by ion-deocclusion and -unbinding. Among these steps, the ion-binding and -unbinding are sensitive to the membrane potential. This is the so-called two-access-channel model (20).

The results and conclusions are significantly different. For the single channel configuration, the pump channel is a pathway connecting the intracellular and extracellular solutions so that Na and K ions are freely moving to cross cell membrane. Orifices at two ends of the channel alternatively open to gate the channel and select ions. For the two-access-channels model, the pumps have two separated channels connected by the “binding pocket” deeply inside the membrane domain where Na and K ions are transported from one access-channel to another by the protein conformational changes through a series of steps including ion-binding, -occlusion in one access-channel and ion-deocclusion and -unbinding in another. It is not clear how the ATP hydrolysis energy released at the cytoplasmic loop drives the conformational changes deeply inside the transmembrane domain.

Which configuration, the single-channel or two-access-channel, is more reliable for the Na/K pumps? Whether Na- and K-ions are carried by the protein conformational changes or freely moving along the channel to cross the cell membrane?

Polytoxin uncouples the extra- and intra-cellular gates (7,8,9), and cysteine mutagenesis introduces cation pathway (10,11,12), neither of which are at physiological structures. In contrast, the dialyzed pump molecules remain the natural physiological structures. Therefore, we chose to re-study the internally dialyzed pumps. In the previous studies, both positive and negative stimulation pulses were applied to the cell membrane and the studies focused on the relaxation charge-movement for both the positive and negative stimulation pulses (16,17,18,19). Here, we only applied a series of negative stimulations in a wide range of magnitude at various ionic concentration gradients. The study is focused on the characteristics of charge movement currents or the relationship between the forward and backward pump currents. We found that the charge-movement pump currents or the obstructed pump channel can only be observed at certain situations. In response to a high stimulation pulse at low ionic concentration gradient, K ions can pass through the channel across the cell membrane. The two-directional charge-movement currents become uni-directional currents indicating that two separated access-channels convert to a single channel configuration. A modified singe-channel configuration model was developed to explain the structures and functions of pump channel.

## Materials of Methods

The experiments were conducted on frog skeletal muscle fibers using the double Vaseline-gap voltage clamp technique. The pump molecules were internally dialyzed by eliminating both the internal and external Na ions. The recipes of the internal and external solutions are as follows.

Na-Free Internal Solution: L-Glutamic acid potassium salt monohydrate (K-glutamate) 58mM; MgSO_4_ 6.8mM; 3-(N-Morpholino) propanesulfonic acid, 4-Morpholinepropanesulfonic acid (MOPS) 5mM; Ethylene glycol-bis (2-aminoethylether)- N,N,N′,N′-tetraacetic acid (EGTA) 20mM; Dibasic potassium phosphate (K_2_HPO_4_) 4mM Cesium hydroxide hydrate (CsOH) 10mM; Adenosine 5′-triphosphate dipotassium salt hydrate (5’-ATP-K_2_) 5mM; Adjust the final pH to 7.30 via KOH at room temperature; store in the freezer at −20 °C.

Na free External Solution: 3,4-Diaminopyridine(3,4-DAP) 3.5mM; Tetramethylammonium chloride (TMA-Cl) 85.35mM; CsCl 10mM; KCl (From 4mM to 36mM); Dibasic potassium phosphate (K_2_HPO_4_) 1.5mM; Potassium dihydrogen phosphate (KH_2_PO_4_) 1mM; CaCl_2_ 1.8mM; BaCl_2_ 1.5mM; Tetrodotoxin (TTX) 1μM; Adjust final pH to 7.10 via HCl at room temperature.

## Results

Figure 1 shows the procedure to obtain the pump currents in responding to a 2 ms, −160 mV stimulation pulse at the extracellular K concentration of 8 mM. Traces a and b were obtained before the application of 300 μM ouabain and traces c and d obtained after as shown in the upper panel. Subtractions of trace b from trace a, and d from c, both showed zero current as shown in the middle panel. Lower panel is the trace c subtracted from trace b, or the ouabain-sensitive pump current. The rising- and falling-phases of the stimulation pulse generate negative and positive transient pump currents, respectively, with the similar magnitude and time-course. This is a typical charge-movement current consisting of forward and backward pump currents. Analyses of the relaxation time-course of the transient pump currents exhibit two components as shown in the lower panel. One is faster with high magnitude and another is slower having lower magnitude. The results are consistent with those obtained previously from the study of axon (Figure 2 in ref.19).

**Figure 1.**
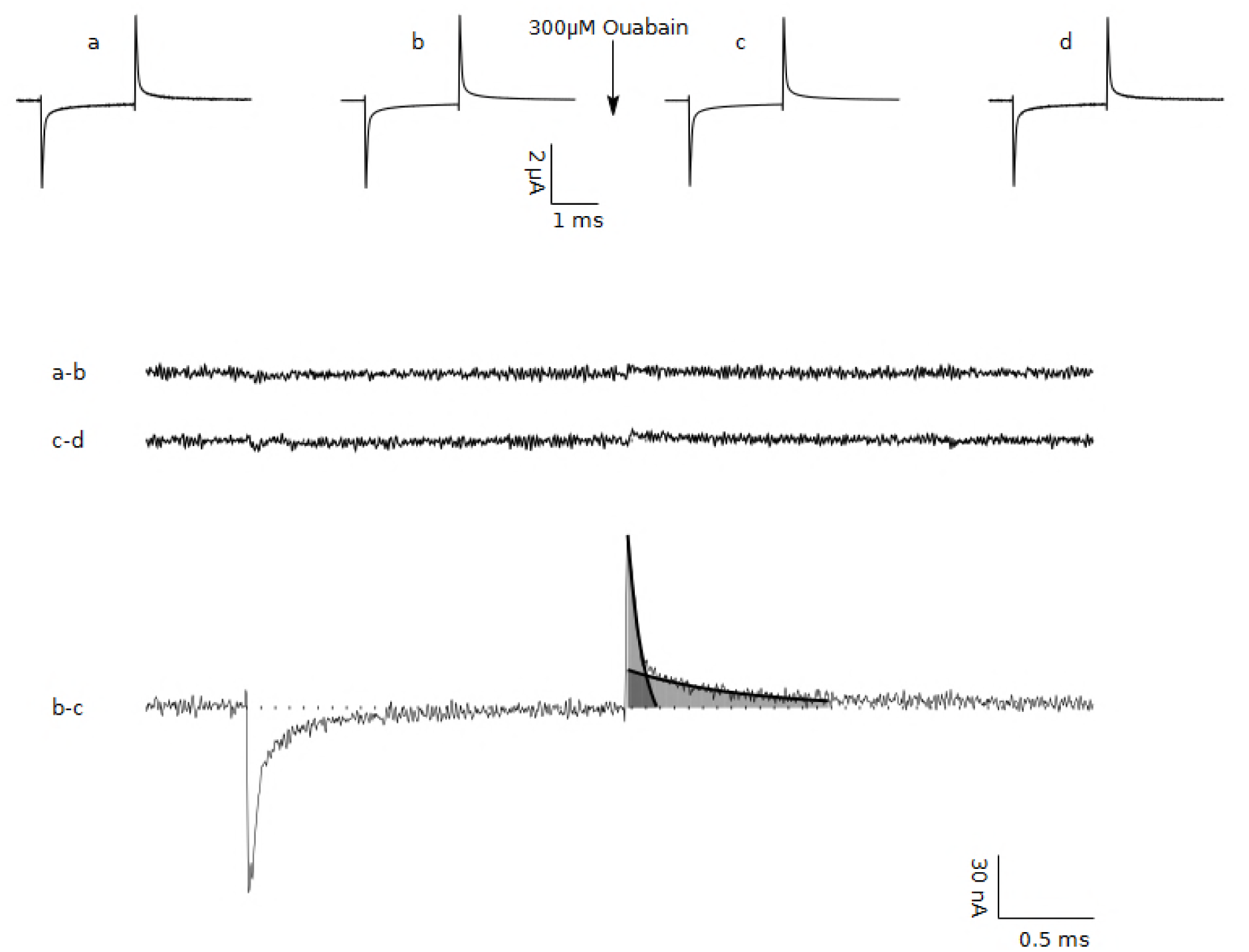
Identification of Na/K pump currents. Upper panel: the transmembrane currents measured before (a and b) and after the application of 300 mM ouabain (c and d) elicited by a 2 ms, −160 mV stimulation pulse. Middle panel: subtraction of trace b from a; and d from c. Lower panel: the Na/K pump currents, or the current c in the present of ouabain from the current b in the absence of ouabain. The backward pump current shows two components of the relaxation time-courses.

**Figure 2.**
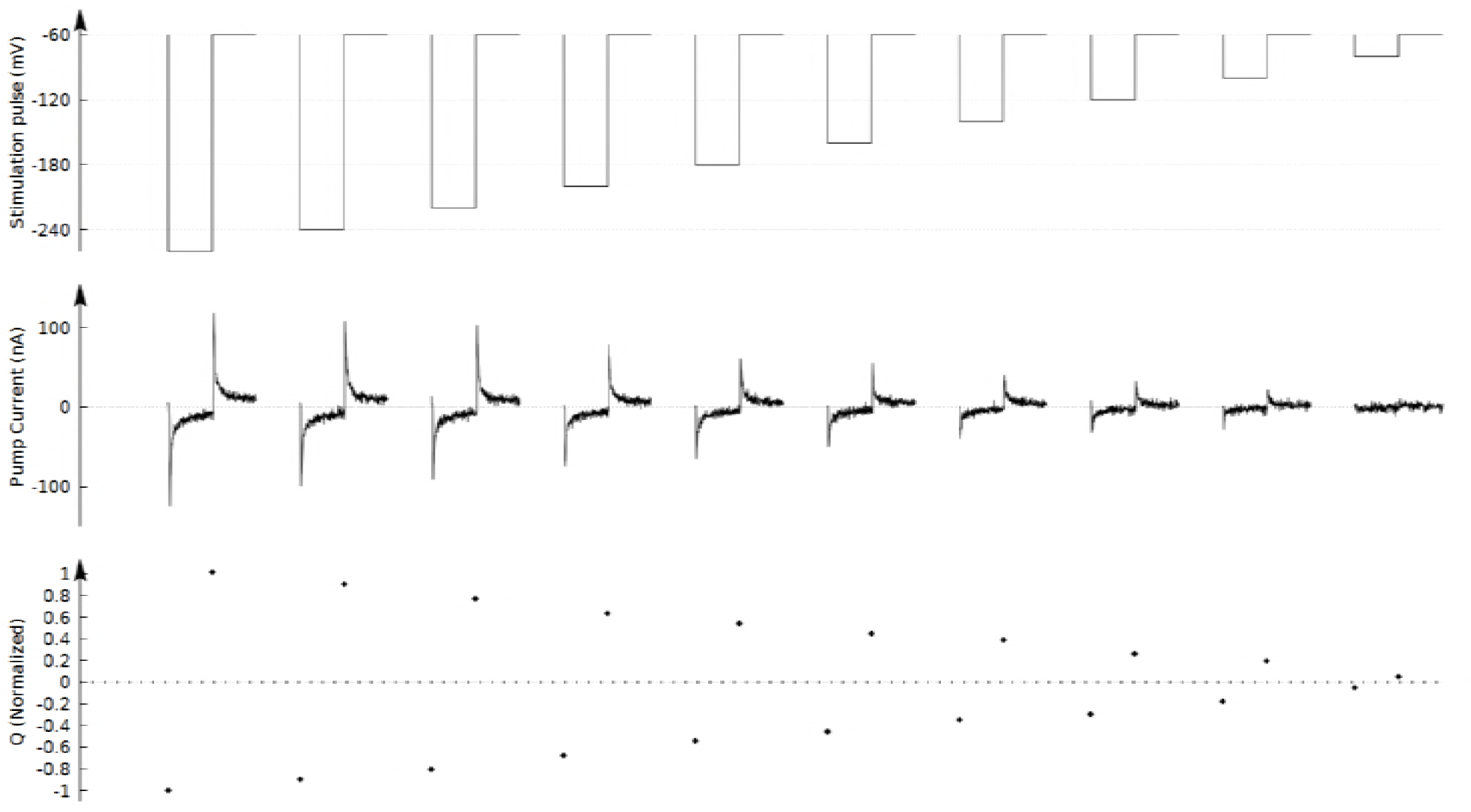
Measurements of the pump currents. Transient pump currents elicited by a series of negative stimulation pulses from −20 to −200 mV. Upper panel is the stimulation pulses. Middle panel: the elicited transient pump currents show the similar magnitude of the forward and backward components. Lower panel: The On-charges and the Off-charges of the transient pump currents have the similar magnitude, indicating the charge-movement-like pump currents

However, in that study (19), the fast component of the off-charge responding to the falling-phase of the stimulation pulse is not considered as the pump currents but some kind of ouabain-induced current. The evidence was that in response to a series of stimulation pulses with the duration of 40, 150, 400 and 1000 μs in measurement of the pump currents, magnitude of the fast component for the off-charge currents remains unchanged (Figure 3 in ref. 19). It is understandable that authors intended to use different durations of the stimulation pulse to dissect the slow and fast components. However, based on the lower panel of the Supplementary Figure 2 (in ref. 19), duration of the fast component is less than 20 μs much short than the narrowest stimulation pulse of 40 μs. It is impossible for a 40 μs or longer stimulation pulse to dissect 20 μs current.

**Figure 3.**
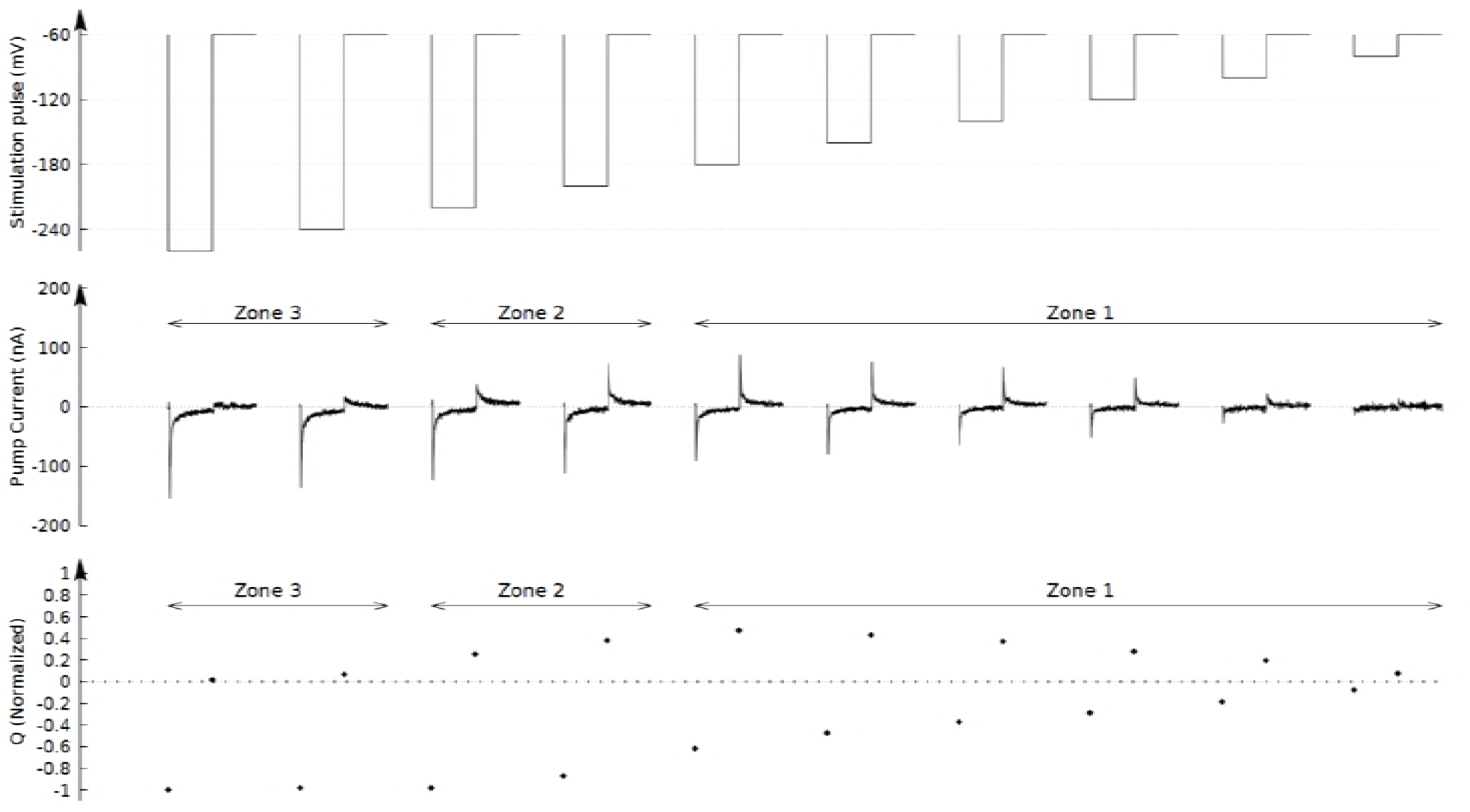
Pump currents measured at 40 mM extracellular K ions. Transient pump currents elicited by a series of negative stimulation pulses at the extracellular K concentration of 40 mM. Upper panel: stimulation pulses; Middle panel: transient pump currents; and Lower panel: On- and Off-charges. Zone 1, responding to the smaller pulses, magnitude of the forward and backward transient pump currents as well as the corresponding charge particles are very similar. Zone 2, for the middle stimulation pulse, the fast components of the backward pump current disappeared while the slow components remain the same. Zone 3, the full backward transient pump currents as well as the Off-charge particle disappear indicating that the K ions pass through the channel reaching the intracellular side.

In addition, as mentioned in the paper, “the fast component has similar time course and magnitude at the On and Off (19).” It is difficult to consider the fast component of the On-charge as the pump current but rule out the fast component of the Off-charge. If the fast component of the off-charge is not the pump currents, the Off-charge will be less than the On-charge particles, which will not satisfy the requirement for the charge-movement current. Moreover, H_2_DTG has long been used to identify Na/K pump current where H_2_DTG-sensitive current has widely accepted as Na/K pump current.

In fact, similar studies of the internally dialyzed pump molecules in the presence of Na ions have showed that the H_2_DTG-sensitive off-charge currents have three components. All three components including fast component have been considered as the Na/K pump currents (18).

We applied a wide range negative stimulation pulses to the cell membrane and studied the relationship between the forward and backward pump currents. Figure 2 shows the pump currents generated by a series of negative stimulation pulses from −20 mV to −200 mV when the cell membrane is held at −60 mV. Indeed, the pump currents are the charge-movement currents having the transient forward and backward components with the similar magnitude. Integration of the transient pump currents for both forward and backward components results in the similar charge particles as shown in lower panel of Figure 2. The higher the magnitude of the stimulation pulses, the larger the transient pump currents and the more the ions were involved, which is the same as the previous study (19).

The charge-movement-like pump currents from the dialyzed Na/K pumps are the main evidence for the two-access-channel model. The transient forward pump current elicited by the rising-phase of the stimulation pulses is always followed by a transient backward pump current generated by the falling-phase of the stimulation pulse. Both the current magnitude and the amount of charge particles are similar, indicating that the K ions moving into the channel driven by the stimulation pulse cannot reach the intracellular side but fully flow back once the stimulation is over. The pump channel is somehow obstructed deeply inside the membrane domain.

However, the similarity of the forward and backward transient pump currents remains only for certain situations. The results changed significantly when we increased the extracellular K concentrations to 40 mM. In response to the stimulation pulses up to −120 mV hyperpolarizing the membrane potential to −180 mV, the forward and backward transient pump currents remain similar, as shown zone 1 in the middle panel of Figure 3. The On-charges remains similar as the Off-charges, as shown in the lower panel.

When the stimulation pulse increased; the backward pump currents elicited by the falling-phase of the stimulation pulses become smaller than the forward pump currents generated by the rising-phase, as shown in zone 2 of Figure 3. Similarly, the Off-charge particles are also smaller than the On-charge particles shown in the lower panel. This indicates that the flowing-in K ions are more than the flowing-out K ions.

More specifically, the fast components of the backward pump currents were smaller than that of the forward pump currents while the slow components remained the same. The larger the stimulation pulse, the less the fast component of the backward pump current is, but the slow component remains the same.

At the magnitude of negative stimulate pulse up to about 160 mV hyperpolarizing the membrane potential to −220 mV, the fast components of the backward transient pump currents fully disappeared, while the slow components had little change. This indicates that the K ions fast flowing into the channel never flow back to the external solution when the stimulation is over. In other words, the fast components of K pump currents become uni-direction currents.

If magnitude of the stimulation pulses further increases, the slow components of the backward pump currents start to reduce. In response to a −200 mV stimulation pulse hyperpolarizing the membrane to −260 mV, the backward pump current with both fast and slow components fully vanished as shown in zone 3. This implies that all the K ions driven by the stimulation pulse flowing into the channel were no longer flowing back when the stimulation is over. The charge-movement pump currents become full uni-directional pump currents.

In other words, at higher extracellular K concentration or lower concentration gradient, a high stimulation pulse can drive K ions to overcome the obstacle in the channel to reach the intracellular solution. The uni-directional single transient pump current indicates a single electrogenic step is involved in the ion-transport across cell membrane.

We also measured the pump currents generated by the negative stimulation pulses at different extracellular K concentrations. Figure 4 shows the ratio of the Off-charge over the On-charge of the pump currents as a function of the magnitude of stimulation pulses at different extracellular K concentrations. The results are averaged from seven experiments. Error bars represent standard deviations. In response to the lower stimulation pulses, the ratio remains at about 1 regardless of the extracellular K concentration, which is the same as shown in zone 1 of Figure 3. The pump currents are the two-directional charge-movement currents. Whatever the K ions flowing into the channel in responding to the stimulation pulse will fully flow back once the stimulation is over.

**Figure 4.**
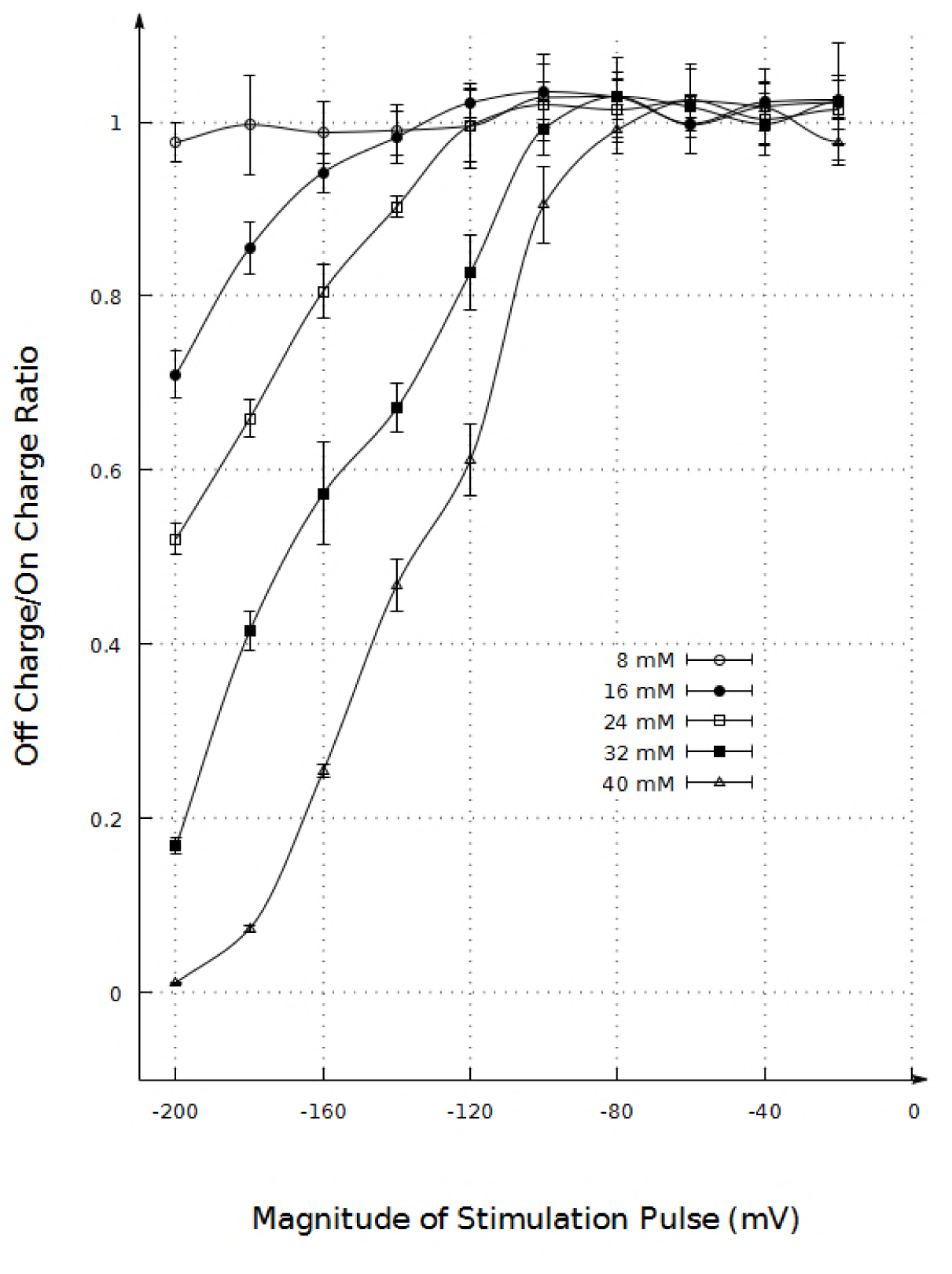
Charge-ratio of the Pump currents at different extracellular K concentrations. Ratio of the Off-charges over the On-charges as a function of extracellular K concentration and the magnitude of stimulation pulses. For the small stimulation pulses up to −100 mV, the ratio remains about 1 regardless of the extracellular K concentrations. The higher the extracellular K ions, the lower the ionic concentration gradient, the smaller the stimulation pulse is required to reduce the ratio. At 40 mM extracellular K, a −200 mV stimulation pulse can fully eliminate the Off-charges.

When the magnitude of stimulation pulse increases, the ratio becomes less than one indicating some of the K ions flowing into the channel no longer flow back, as shown in zone 2 in Figure 3. The higher the concentration of the extracellular K ions, or the lower the ionic concentration gradient, the less the ratio is, or the easier for the K ions moving through the channel into the intracellular solution. At about 40 Mm extracellular K ions, in response to a −200 mV stimulation pulse, the ratio is close to zero, equivalent to zone 3 of Figure 3. The pump currents are no longer the two-directional charge-movement currents but become uni-directional currents, K ions passing through the channel across the cell membrane.

These results are difficult to explain with the two-access-channel model. First, because the pumps are dialyzed, no ATP hydrolysis energy is available to drive the protein conformational changes. It is impossible for an electric field to generate the protein conformational changes through a series of steps. Then, how can the K ions being transported from one access-channel to another showing a uni-directional pump current?

Second, because both the ion-binding and -unbinding steps in two access-channels are sensitive to the membrane potential resulting in two separated electrogenic steps, the pump currents crossing cell membrane should exhibit two transient components. Why do the pump currents only have one transient component?

Third, we will show latter that duration of the pump currents across cell membrane is only about 100 μs, which is much shorter than both the channel currents (in ms) and gating currents (hundreds of μs) for Na channels. Na channel is the protein in living system with the fastest electrical response this is because of involvement of generation and propagation of action potential. The conformational changes are relatively simple only involving shifting of a few charged amino acids in S4 domain to open or close the gate (21). It is difficult for the pump molecules in such short time to complete a series of more complicated protein conformational changes to transport ions from one access-channel to another.

### A modified single-channel model

Here, we present a model of the modified single-channel configuration. First, comparison of the two-directional charge-movements currents (either positive and negative) shown in Figure 2 and the unidirectional transient pump currents across cell membrane shown in zone 3 of Figure 3, durations of the pump currents are very similar. The former are the K ions moving within the internal access-channel without any conformational changes while the latter are the ions moving further to the intracellular side across the cell membrane. The similar duration indicates that K ions moving across the cell membrane did not involve conformational change. The pump channel has a single-channel configuration.

Second, small stimulation pulse cannot drive K ions moving across cell membrane while a large stimulation can. Considering the ionic concentration gradient against the ion movement, the dependence of the ion-movements on the magnitude of stimulation pulse implies that energy, instead of protein conformational changes, dominates the ion-movement across cell membrane which further confirms the single-channel configuration.

Third, the extremely short time-course of the transient pump currents across the cell membrane indicates that K ions must quickly move along the channel or have high moving speed. This can also be observed by the high magnitude of the transient pump currents. The faster the moving speed of the ions, the higher the magnitude of the pump currents is.

Forth, orifices at two ends of the channel have been showed by the cysteine-scanning mutagenesis studies of α-helices in transmembrane domain with MTSET or MTSES (10,11,12). Functions of the orifices are to isolate the channel lumen from the environments, or occlusion/de-occlusion of Na or K ions. Another function of the orifices is to select Na or K ions. The mechanisms involved in the ion-selectivity may be adapted from the ion-channel based on the ion size or ion-dehydration energy (21). Existence of orifices can be partially proved by the limited pump currents generated by the small stimulation pulses shown in Figures 2 and 3. In response to a low stimulation pulse of 20 mV, no pump current can be generated. Ions passing through the orifice inevitably require energy. The 20 mV stimulation pulse may not be high enough to drive K ions passing through the orifice into the channel. In contrast, if it is a wide open access-channel, the transient pump current should be generated regardless of the magnitude of stimulation pulse.

Orifice of the pump channel can also be partially proved by the distribution of the charged amino acids in the α-helices of the transmembrane domain. Upper panel of Figure 5 shows the sequence of amino acids in all ten α-helices in the transmembrane domain (22,23,24). At two ends of the channel there are both negatively (black dot) and positively (white dot) charged amino acids distributed in different α-helices. Electrostatic attracting force among those positively and negatively charged amino acids facilitates the formation of the channel orifice, as shown in the lower panel.

**Figure 5.**
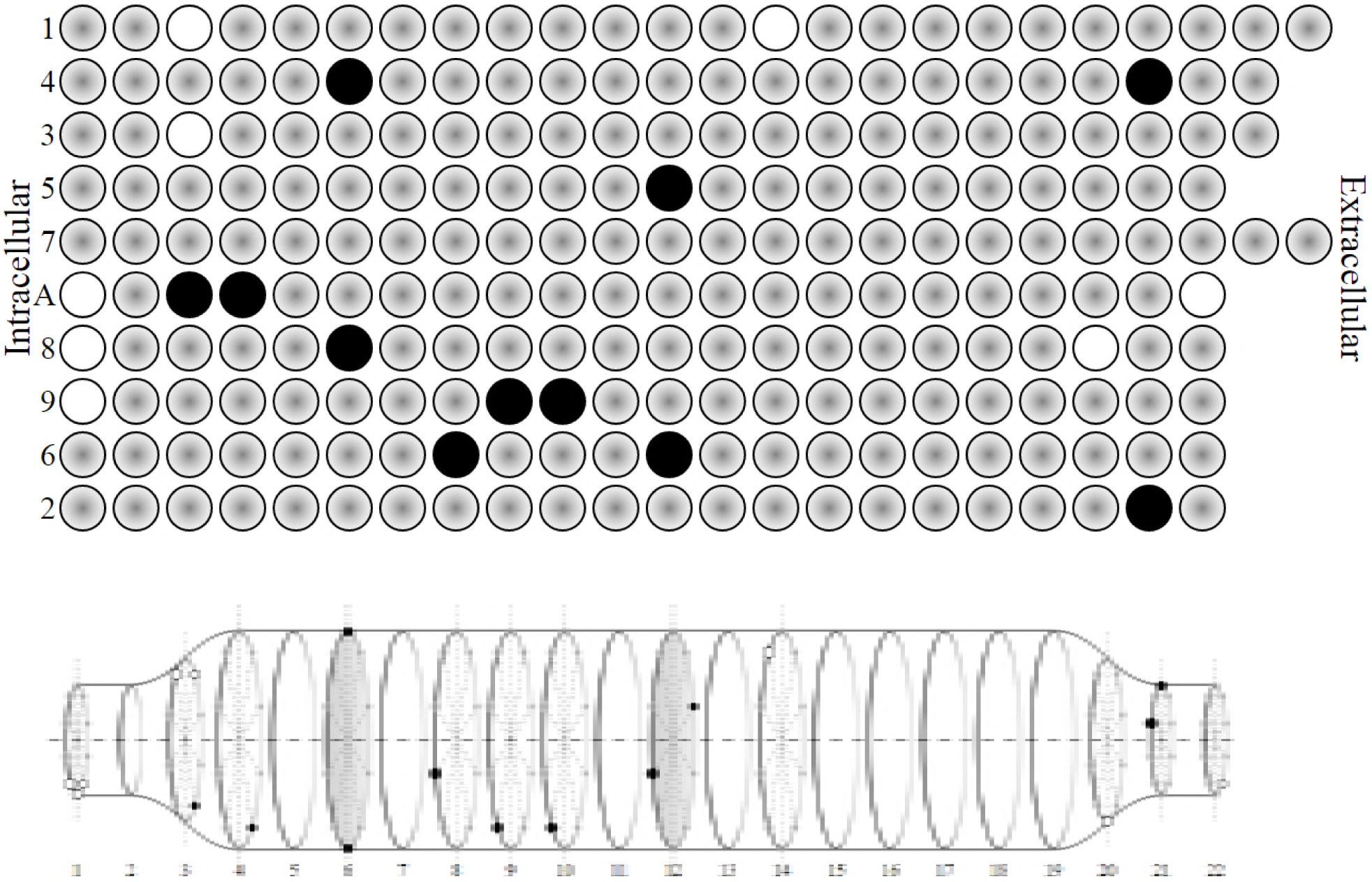
Pump channel structures. Upper panel: the amino acids in all ten α-helices of the transmembrane domain. Black dots represent negatively charged amino acids and white dots represent the positively charged amino acids. Lower panel: pump channel formed by the α-helices in the transmembrane domain. At two ends of the channel, electrostatic attracting force among the negatively and positively charged amino acids facilitates formation of the orifice, while the negatively charged amino acids in the middle of channel form an energy-array in spatial functioning as a collimator to align the moving ions in the lumen of channel.

Indeed, studies of palytoxin-treated pump molecules (7,8,9) and molecular dynamic simulations of the crystallized pump structure (19) have showed the wide open vestibule of the access-channel. The former disrupted the coupling of the intra- and extracellular gate enabling both gates opening simultaneously, which inevitably affect the natural protein structures. The latter is based on the crystallized structures of pump molecules. It is not clear how the crystallized structures reflect the protein structures at physiological running condition.

For example, during the protein purification with the SDS treatment, water molecules may get into the lumen of channel. Then, the water hydrophilic force may push the hydrophobic α-helices to open the orifice of the channel resulting in the wide open vestibule. Interestingly, crystallized structures of K-channel also exhibit wide open vestibule (25). Both the functions and structures for the Na/K pumps and the ion-channels differ significantly. It is not clear why the crystallized structures both show the wide open vestibule.

Fifth, in order to ensure the fast moving ions passing though the channel, pump molecules must have some kind special arrangements in the channel to avoid the moving ions hitting the wall of channel. Interestingly, deeply inside the α-helices of the transmembrane domain there are mainly negatively charged amino acids as shown in upper panel of Figure 5. Those negatively charged amino acids located at different α-helices, about one at each, may form some kinds of energy array in space functioning as a collimator to align the moving ions in the lumen of channel. Molecular simulations show that the two negatively charged amino acids in turn-6 can form the collimator in Y-direction and the two negatively charged amino acids in turn-12 can form a collimator in X-axis, as shown in lower panel of Figure 5.

As a side-effect, the negatively charged amino acids inevitably attract the Na and K ions moving along the channel or form an energy-well for the moving cations. It is necessary to point out the difference between individual negatively charged amino acids and the collimator formed by multiple spatially distributed negatively charged amino acids. For the former, the amino acids attract cations and eventually can bind with them where kinetic energy of ions is lost. In order to unbind the ions, extra energy has to be consumed. For the latter, the collimator only aligns the moving ions in the lumen of channel but can never bind with the cations so that no extra-energy is required. If the initial speed of the cations can overcome the ionic concentration gradient, the collimator will allow the ions passing through. If not, the ions will be stopped in the collimator.

Previously, energy-barrier has been proposed in the channel to explain the obstacle of the ion movements (26,27,28). That is lack of the structural base because of no positively charged amino acids in the α-helices to form an energy-barrier for cations. In addition, from the viewpoint of energy, it is unlikely. Extra energy will be required to overcome the barrier which inevitably reduces the energy efficiency for Na/K pump.

For the dialyzed pumps without ATP hydrolysis energy, if the stimulation pulse is high enough as shown in this paper, K ions can pass through the collimator across cell membrane showing the unidirectional pump currents. However, if the stimulation pulse is not high enough to overcome the ionic concentration gradient, the ions will be stopped in the energy-well, as if the channel is ended deeply inside the membrane showing transient forward pump currents. Once the stimulation is over, the ionic concentration gradient will drive the ions flowing back resulting in transient backward pump currents. At physiological running condition, ATP hydrolysis energy is high enough so that Na and K ions can pass through the collimator across cell membrane against the ionic concentration gradient.

### Using the modified single-channel configuration to explain the experimental results

#### Two components of the transient pump currents

Previously, the three components of the transient Na currents have been interpreted as the discrete, sequential steps in the ion-binding and -unbinding with the binding-site (18). In fact, the sequential pattern can also be obtained from the modified single-channel configuration. In order to select Na or K ions into the channel, ions have to pass the orifice one-by-one in a sequential pattern. In addition, once getting into the channel, the electrostatic force among two ions will inevitably repel each other. The ion first getting into the channel will move faster result in the fast component of the transient pump currents with a higher magnitude, while the ion getting-in later will move slower result in the slow component with lower magnitude. If the electric stimulation pulse is not high enough to overcome the ionic concentration gradient, 2 K ions will be stopped in the collimator in a sequential pattern. Once the stimulation is over, with the same token, 2 K ions will be driven back by the ionic concentration gradient sequentially out of the channel. Upper panel of Figure 6 shows the movement of two K ions. High bar with narrow duration represents the movement of the first K ion followed by the low bar with wide duration representing the second K-movement.

**Figure 6.**
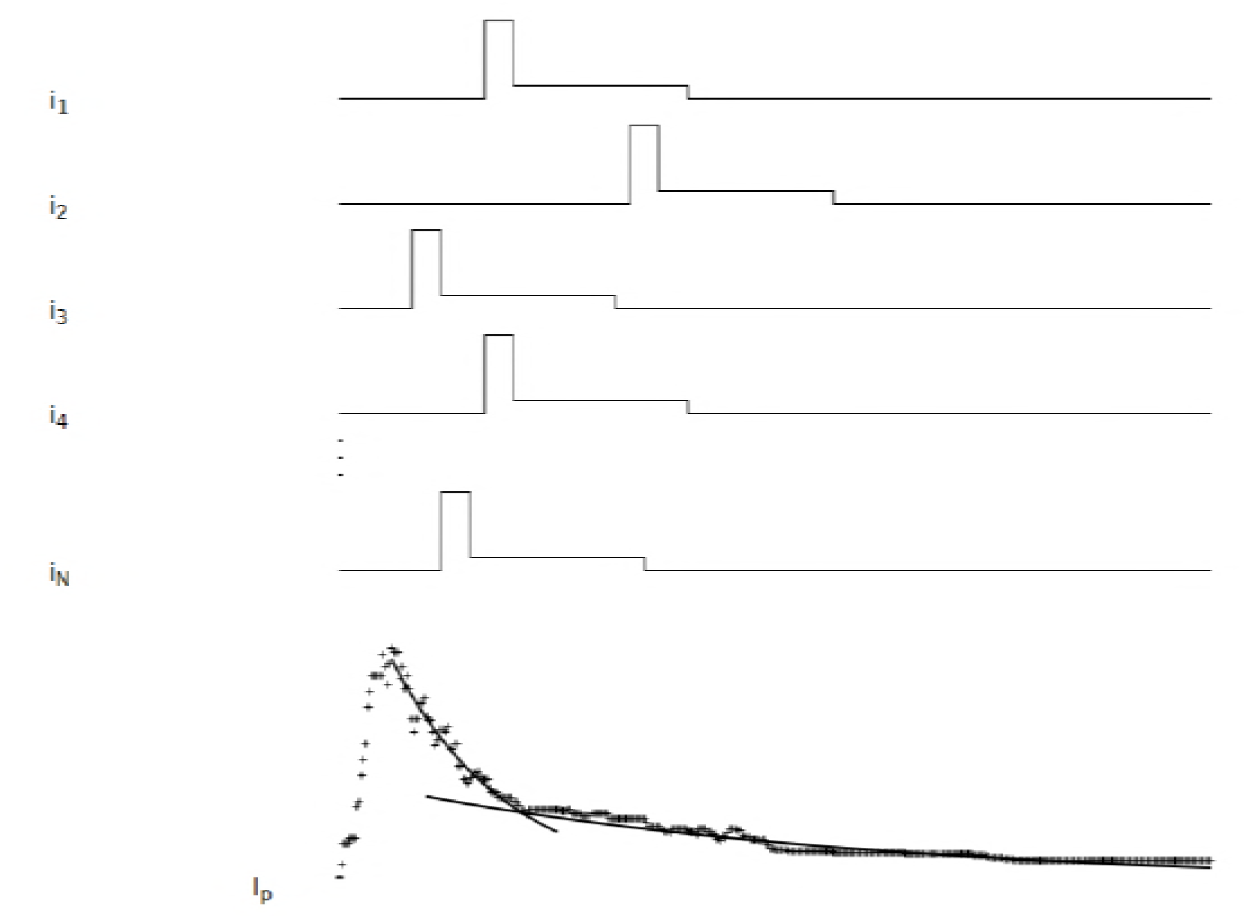
Two exponential decays of the pump currents. Upper traces represent the single pump currents distributed statistically. Each pump current consists of two bars. The high bars with short duration represent the first K ions getting into the channel moving fast while the low bars with long duration stand for the second K ions getting-in the channel moving slower. Bottom trace: the macroscopic pump currents from the current-summation (N=200) showing two components of the transient pump currents. Two solid lines are the fitted curves for the two components.

Because a large amount Na/K pumps is involved in the fibers, the measured pump currents is a summation from all the pumps which may have some distribution due to imperfect synchronization. As a result, the total transient pump currents exhibit two exponential decays. The fast component represents the current summation for the first fast moving K ions from all the pumps, while the slow component stands for the current summation from all the second slow moving K ions, as shown in the lower panel of Figure 6. This technique has been used to explain the shape of Na-channel currents from the whole-cell voltage-clamp experiments by the currents from the single-channel patch clamp measurements (29,30).

#### Reduction and elimination of the backward pump currents for the dialyzed pumps

It is necessary to point out for the dialyzed pumps, no ATP hydrolysis energy is available. Electric field is the only driving force for the ion-movements to overcome the ionic concentration gradients. This can be seen by observing the relationship of the rising-phase of the membrane potential, the capacitance currents, as well as the transient pump currents shown in Figure 7. Please notice the different scale for the currents. The total transmembrane current (mainly the capacitance currents) is associated with the rising-phase of the membrane potential (middle panel) because capacitance current is proportional to the first directive of the membrane potential. In contrast, the pump currents are delayed. Membrane potential has to be fully established before the transient pump currents start to increase and the peak is postponed for tens of μs. This is similar as the voltage-dependent Na and K channel currents where some K-channels are even called delayed rectifier K channel (21).

**Figure 7.**
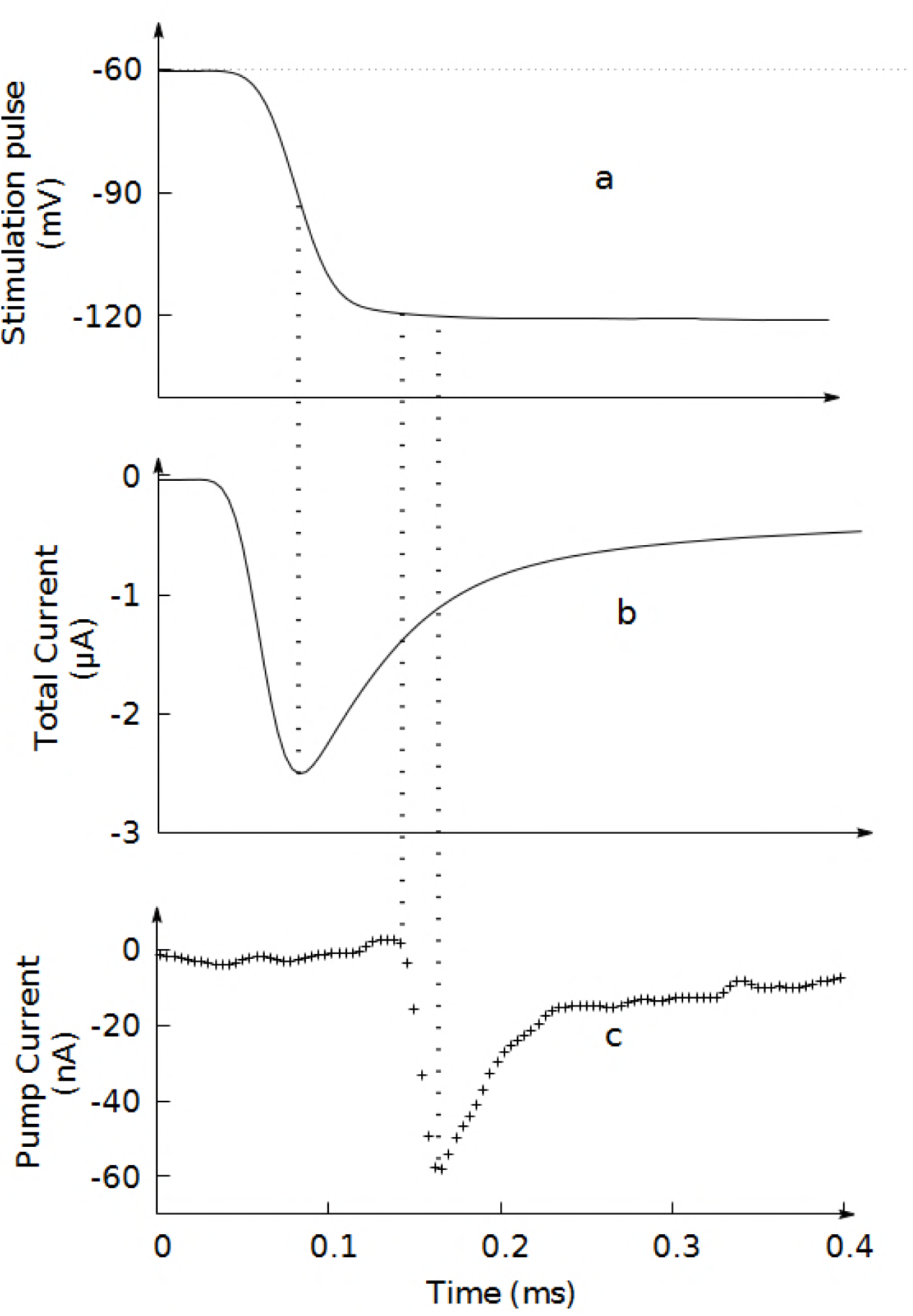
Comparison of the membrane potential and the transient pump current. Upper panel: Rising-phase of the membrane potential; Middle panel: the capacitance currents; Lower panel: the transient pump currents. Pump currents start to increase after the membrane potential has been fully established and the peak values delays for a few tens of μs.

Trace a in Figure 8 is a cartoon of the modified single-channel configuration with the collimator or energy-well. Trace b represents the intrinsic energy-profile along the pump channel with the energy-well in the middle of channel. At 40 mM extracellular K concentration, the K-equilibrium potential is about −35 mV. The K moving-in against the concentration gradient can be expressed as the overcome of the equilibrium potential difference, the intracellular positive with respective to the reference of 0 mV at the extracellular side. Therefore, the energy-profile is titled to the positive direction with respect to the extracellular side, as shown in the trace c of Figure 8. When the cell membrane was held at a holding potential of −60 mV, the energy-profile tilts back a little but still to the positive direction, as shown in Trace d of Figure 8. This is because the majority of the applied membrane potential is dropped on the protein structure and little is left in the channel space to drive the ion-movements.

**Figure 8.**
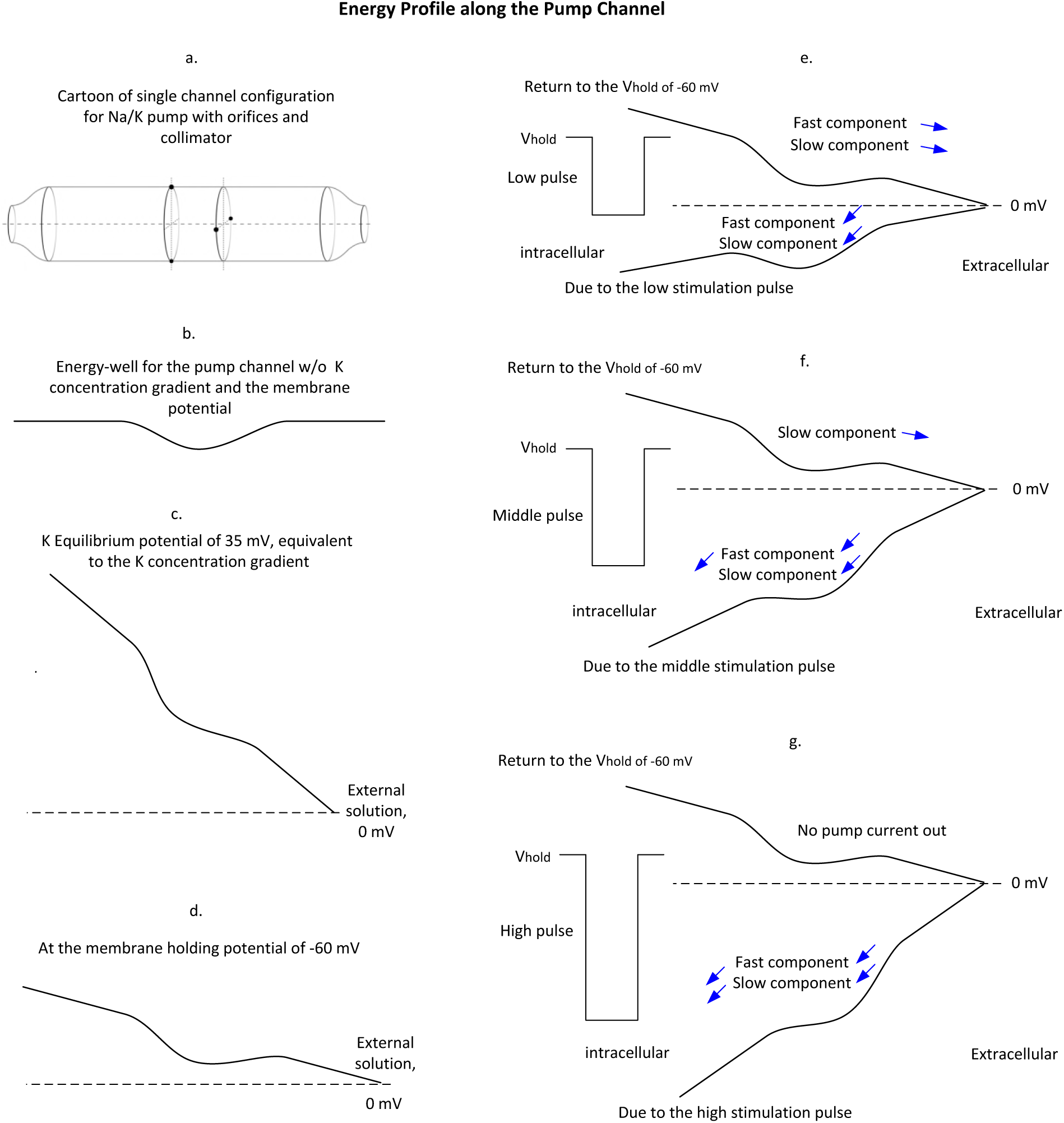
Interpretation of the dialyzed pump currents. Trace a: Cartoon of the pump channel with a collimator inside the channel. Trace b: The intrinsic energy-profile of the pump channel with energy-well in the middle of channel. Trace c: the energy-profile tilts to the positive direction due to the ionic concentration gradient (40 mM extracellular K ions) expressed as the K-equilibrium potential of 35 mV (intracellular positive). Trace d: the energy-profile tilts back a little due to the membrane potential is hold at −60 mV. Trace e: Energy-profile changes responding to a small negative stimulation pulse. Trace f: Energy-profile changes due to the application of a middle stimulation pulse. Trace g: Energy-profile changes in response to a high magnitude stimulation pulse. See texts.

In responding to a negative stimulation pulse with a low magnitude, such as −80 mV hyperpolarizing the membrane potential to −140 mV, the energy-profile of the pump channel is tilted to the negative direction as shown in trace e. The K ions, once getting into the channel, will be driven by the stimulation pulse and attracted by the energy-well. Due to the low pulse magnitude, the electric field cannot overcome the ionic concentration gradient. Even though the energy-well attracts the K ions resulting in the transient inward pump currents, K ions will be eventually stopped in the energy-well as if reaching the end of channel. Once, the stimulation is over, cell membrane is returned to the holding potential, the energy-profile tilts back to the positive position so that the ionic concentration gradient will drive all the inward K ions out of the channel resulting in the outward transient pump currents, as shown in trace e in Figure 8. The results are also shown in zone 1 of Figure 3.

The higher the magnitude of the negative stimulation pulse, the more the negative the energy-profile tilts to. In response to a medium stimulation pulse, such as - 160 mV hyperpolarizing the membrane potential to −220 mV, the energy carried by the fast components of the inward K ions will be able to overcome the ionic concentration gradient passing through the collimator flowing into the intracellular solution, while the slow component cannot because of its lower energy. As the stimulation pulse is removed, only the slow component flows back resulting in the smaller backward pump currents than the forward pump currents as shown in zone 2 of Figure 3 and trace f in Figure 8.

In fact, the smaller fast components of the transient backward pump currents than that in the forward pump currents in responding to a −110 mV stimulation pulse have been observed previously in study of the release of Na ions from the internally dialyzed pump molecules (18). It was called no-conservation of the fast charges between on and off voltage steps. However, the involved underlying mechanism was not further studied.

When the stimulation pulse has high magnitude, such as −200 mV hyperpolarizing the membrane potential to −260 mV, the energy-profile will be further tilted to the negative direction. All the K ions including the slow component will be more than enough to overcome the ionic concentration gradient and flowing into the intracellular solution. Therefore, there is no backward pump current in responding to the falling phase of the stimulation pulse, as shown zone 3 in Figure 3 and trace g in Figure This indicates that the obstacle in the channel can no longer hinder the ion movements. The two access-channels become a single channel configuration.

When the intracellular K concentration remains unchanged, the higher the extracellular K concentration, the less the concentration gradient, the lower the equilibrium potential of K ions, and therefore the less the energy-profile tilted to the positive direction. Consequently, in responding to the same negative stimulation pulse, the more the energy profile tilted to the negative direction, or the more the K ions can pass the energy-well flowing into the intracellular solution. In other words, the higher the extracellular K concentration, the less the magnitude of the stimulation pulse is required to drive K ions in overcoming the ionic concentration gradient to demonstrate the single channel configuration.

### Effect of the extracellular pH value on the pump currents

In order to confirm the effects of collimator and energy-well due to the negatively charged amino acids deeply inside the pump channel, we redo the above experiments in the extremely low pH value, 4.6 of the external solution. The extracellular K concentration was remained at 40 mM, and again, the negative stimulation pulses from −20 mV up to −200 mV were applied to the cell membrane. The measured pump currents were shown in Figure 9. Interestingly, in response to the same magnitude of stimulation pulses, the transient pump currents were significantly reduced but the plateau increased. These results are similar as those obtained previously from other labs (31,32). Most impressively, regardless of the increments in the stimulation pulse, the transient backward pump currents responding to the falling phase only slightly decreased but never disappeared. This demonstrates that most of the flowing-in K ions driven by the stimulation pulse cannot pass through the channel across the cell membrane but returning to the external solution once the stimulation is over.

**Figure 9.**
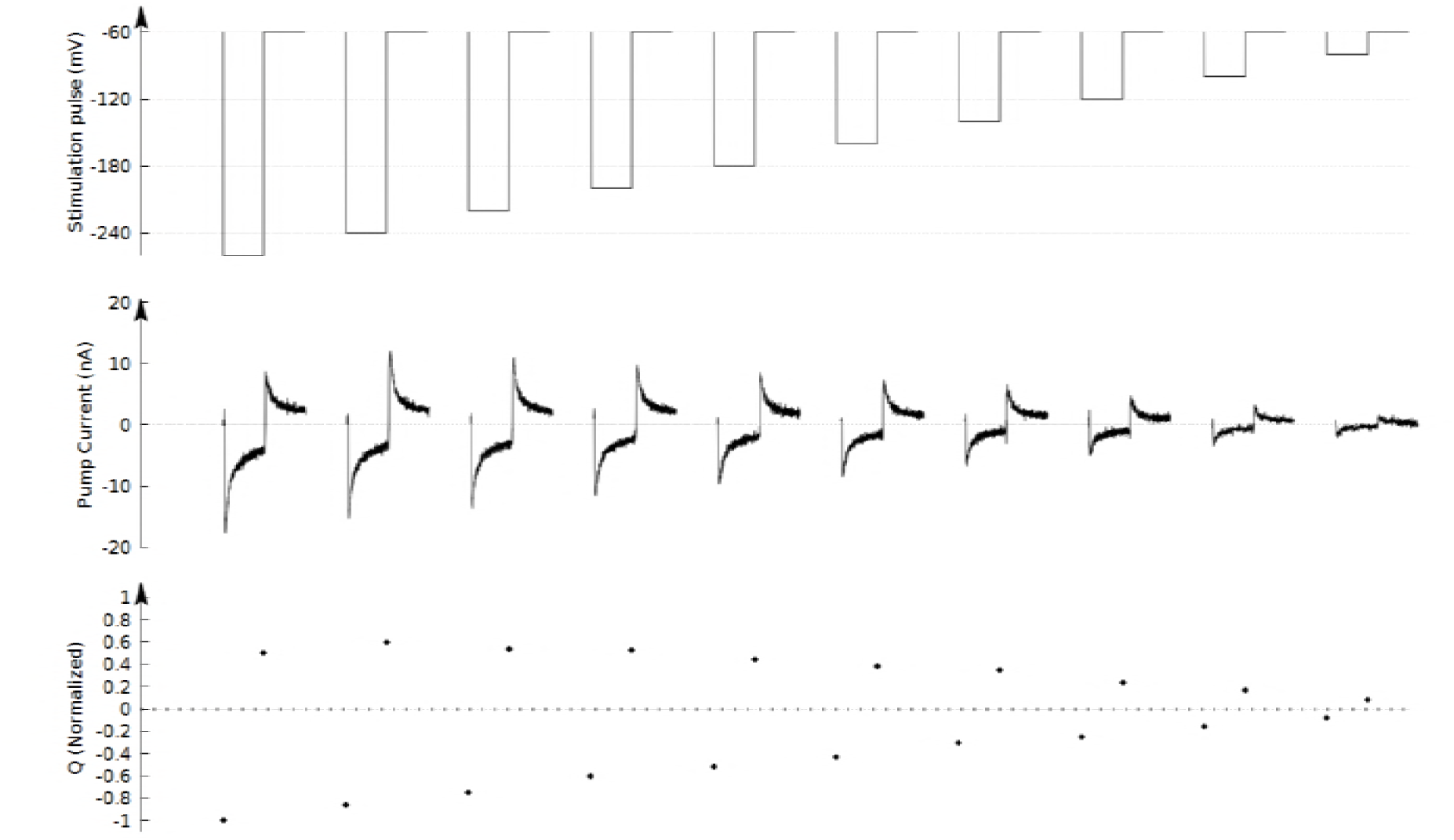
low pH value affects the pump currents. Upper panel: a series of stimulation pulses from −20 to −200 mV at 40 mM extracellular K concentration. Middle panel: the elicited transient pump currents. Lower panel: the corresponding On-charges and the Off-charges. Due to low pH value, magnitude of the transient pump currents significantly reduced while the plateau increased. Regardless of the magnitude of stimulation pulses, the backward transient pump currents responding to the falling phase of the stimulation pulse never disappeared.

Comparing pH values of 4.6 and 7.1 at physiological condition, concentration of hydrogen ions increased almost by three orders. Hydrogen ions smaller than either Na or K ions can pass through the orifice into the channel and interact with the negatively charged amino acids. Due to neutralization of the negatively charged amino acids, there will be less or no energy-well to attract the flowing-in K ions. The only driving force for the moving ions is the applied stimulation pulse. The lower speed of the flowing-in K ions result in the lower magnitude of the transient pump currents and the higher plateau. In addition, because the collimator functions is reduced. The flowing-in K ions cannot be well aligned in the lumen of pump channel. Consequently, except smaller amount of K ions can pass through the channel reaching the intracellular side, most of them may hit the wall of channel being reflected back and forth. Therefore, regardless of the magnitude of stimulation pulse, most of the flowing-in K ions will flow back once the stimulation is over.

## Discussion

In this study, we re-investigated the internally dialyzed Na/K pumps, the experimental foundation for the two-access-channel model. We found that the charge movement pump currents, the main evidence of the two-access-channels model, are only observed in response to relatively small stimulations. For larger stimulations at lower K concentration gradient, the backward pump currents disappeared. The two-directional charge-movement pump currents become uni-direction currents indicating that K ions pass through the channel across the cell membrane, or, the two access-channels become a single channel configuration.

The pulse magnitude-dependent behavior of the pump channel indicates that it is energy, instead of the protein conformational changes, dominating the ion-transport across the cell membrane.

A model of modified single-channel configuration is introduced which well explains the experimental results. Na and K ions are freely moving along the pump channel. The negatively charged amino acids deeply inside the channel form a spatial energy-array functioning as a collimator and energy-well for the moving cations. For the dialyzed pumps without ATP hydrolysis energy, a small stimulation pulse cannot drive K ions to overcome the ionic concentration gradient so that the K ions will be stopped in the collimator or energy-well. Once the stimulation is over, ionic concentration gradient will drive K ions back to the external solution. The results imply that as if the pump channel is obstructed into two segments or two access-channels. However, if the stimulation pulse is high enough that can drive K ions to overcome the ionic concentration gradient, the ions can pass through the channel across the cell membrane showing a single-channel configuration.

### Dialyzed pumps vs. synchronized pumps at physiological running condition

Due to random pumping pace, study of a group of pumps at physiological running condition cannot reveal the details of pump molecules. For the dialyzed pumps, pumping cycle from all the pumps are interrupted at the same step. Stimulation triggers the partial reaction from all the pumps simultaneously. From this aspect, the dialyzed pump is a platform to study details of Na/K pumps. However, the pumps are only synchronized to the specific step where the pumping cycle was interrupted. Because of the inability to hydrolyze ATP, the electric stimulation-triggered partial reactions cannot reflect the pump functions at physiological condition.

For example, as mentioned above, the ionic concentration gradient for the K ions is only about 35 mV of the equilibrium potential. Nevertheless, we have to use an extremely large stimulation pulse of −200 mV at the membrane holding potential of −60 mV, totally −260 mV, to overcome the ionic concentration of 35 mV in order to drive K ions across the cell membrane. Otherwise, K ions cannot pass through the channel. As mentioned above, large amount of the applied electric field is dropped on the protein structure, and little is left in the channel to drive ion-movements.

At physiological running condition ATP hydrolysis energy can directly drive Na and K ions moving along the channel to overcome the ionic concentration gradient regardless of the membrane potential. Therefore, the pumps always show a single-channel configuration in a wide range of the membrane potentials.

As a result, in order to study the details of Na/K pumps, we need to synchronize the pump molecules at physiological running condition. Once the pumps are synchronized to all the steps throughout the pump cycle, signals from the individual steps will be superimposed correspondingly. The synchronized pump currents will behavior like a real-time semi single-pump current revealing many details of the Na/K pumps. The results will be reported separately.

## Acknowledgement

The project is partially supported by NIH grant 2R01 50785 (W.C.) and NSF grant 0515787(W.C.)

Liang PF conducted experiments in study of the pump currents; Mast J performed the molecular dynamics simulations for the pump channel; Chen W developed modified single channel configuration, wrote the paper, and supervise the project.

## Author Information

authors declare no competing financial interests.

## References

1. Lauger P., A channel mechanism for electrogenic ion pumps, Biochim. Biophys Acta 552:143–161 (1979).

2. Gadsby, D.C., Rakowski, R.F., & De Weer, P., Extracellular access to the Na,K pump: pathway similar to ion channel. Science 260: 100–103 (1992).

3. Hilgemann, D.W., Channel-like function of the Na, K pump probed at microsecond resolution in giant membrane patches, Science, 263:1429–1432 (1994).

4. Reyes, N., and Gadsby, D.C., Ion permeation through the Na, K-ATPase, Nature, 443(470-474), 2006.

5. Takeuchi, A., Reyes, N., Artigas, P., & Gadsby, D.C., The ion pathway through the opened Na,K-ATPase pump, Nature, 456:416–423 (2008).

6. Gadsby, D.C., Ion channels versus ion pumps: the principal difference, in principle, Nature Review 10:344–352 (2009).

7. Kim, S.Y., Marx, K.A., & Wu, C.H., Involvement of the Na, K-ATPase in the induction of ion channels by palytoxin, Naunyn-Schmiedeberg’s Arch Pharmacol, 351:542–554 (1994).

8. Artigas, P., & Gadsby, D.C., Na/K pump ligands modulate gating palytoxin-induced ion channels, Proc, Natl, Acad, Sci, USA, 100:501–505 (2003).

9. Artigas, P., & Gadsby, D.C., Large diameter of palytoxin-induced Na/K pump channels and modulation of palytoxin interaction by Na/K pump ligands, J. Gen, Physiol, 123:357–376 (2004).

10. Horisberger JD, Kharoubi-Hess S, Guennoun S, and Michielin O, 2004, The fourth transmembrane segment of the Na, K-ATPase a subunit, The Journal of Biological Chemistry, 279(28):29542–29550.

11. Guennoun S, Horisberger JD, 2000, Structure of the 5th transmembrane segment of the Na, K-ATPase a subunit: a cysteine-scanning mutagenesis study, FEBS Letters 482:144–148.

12. Guennoun S, Horisberger JD, 2002, Cysteine-scanning mutagenesis study of the sixth transmembrane segment of the Na, K-ATPase a subunit, FEBS 513:277–281.

13. Chen, W., Synchronization of carrier-mediated pump molecules by an oscillating electric field: Theory, Journal of Physical Chemistry B, 112(32), 10064–70 (2008).

14. Chen, W., Zhang, Z.S. & Huang, F., Synchronization of the Na/K pumps by an oscillating electric field, Journal of Bioenergetics and Biomembrane, 40:347–357 (2008).

15. Gadsby, D.C., Ion channels versus ion pumps: the principal difference, in principle, Nature Review 10:344–352 (2009).

16. Holmgren, M., Wagg, H., Bezanilla, F., Rakowski, R.F., De Weer, Gadsby, D.C., Three distinct and sequential steps in the release of sodium ions by the Na/K-ATPase. Nature, V(403): 898–901 (2000).

17. Castillo, J.P. et al, Energy landscape of the reactions governing the Na deeply occluded state of the Na/K ATPase in the gland axon of the Humboldt squid Proc. Natl Acad. Sci. USA 108, 20556-20561, (2011).

18. Gadsby, D.C., Bezanilla, F., Rakowski, R.F., De Weer, P. & Holmgren, M., The dynamic relationships between the three events that release individual Na ions from the Na/K-ATPase, Nature Communication, 3:669 doi:10.1038/ncomms1673 (2012).

19. Castillo, J.P., Rui, H., Basillio, D., Das, A., Roux, B., Latorre, R., Bezanilla, F. & Holmgren, M., Mechanism of potassium ion uptake by the Na/K-ATPase, Nature Communications, 6:7622 doi:10.1038/ncomms8622 (2015).

20. Apell, H.J, How do P-type ATPases transport ions? Bioelectrochemistry, 63:149–156 (2004).

21. Hille B., Ion channel of excitable membranes, 3rd edition, Sinauer Associates Inc (2001).

22. Shinoda, T., Ogawa, H., Cornelius, F., and Toyoshima, C., Crystal structure of the sodium potassium pump at 2.4 A resolution, Nature 459(446-450), 2009.

23. Kanai, R., Ogawa, H., Vilsen, B., Cornelius, F., & Toyoshima, C. Crystal structure of a Na+-bound Na+,K+-ATPase preceding the E1P state. Nature 502, 201–206 (2013).

24. Hanai, R., Ogawa, H., Vilsen, B., Cornelius, F., & Toyoshima, C., Crystal structure of a Na+-bound Na+, K+-ATPase preceding the E1P state, Nature, 502:201–207 (2013).

25. Jogini V, and Roux B, Dynamics of the Kv1.2 voltage-gated K+ channel in a membrane environment, Biophysical Journal, 93:3070–3082.

26. Glynn, I.M., The Na,K-transporting adenosine triphosphatase, in A.N. Martonosi (ed.), The enzymes of biological membrane, Vol 3, 2nd Ed. Plenum Press, New York, pp 35–114 (1985)

27. Forbush III, B, Occluded ions and Na, K-ATPase, In J.C. Skou, J.G. Norby, A.B. Maunsbach and M Esmann (des.) The Na,K-pump, Part A: Molecular aspect. Progr. Clin. Biol. Res. 268A. A.R. Liss, New York, pp 229–248 (1988).

28. Lauger, P., Electrogenic ion pumps, Sinauer Associates, Inc, (1991).

29. Sigworth, R., and Neher, E., Single Na channel currents observed in cultured rat muscle cells, Nature, 287:447–449, (1980).

30. Patlak, J., and Horn, R., Effect of N-bromoacetamide on single sodium channel currents in excised membrane patches, J. Gen. Physiol., 79:333–351(1982)

31. Salonikidis P.S., Kirichenko S.N., Tatjanenko L.V., Schwarz W., and Vasilets L.A., Extracellular pH modulates kinetics of the Na, K-ATPase, Biochimica et Biophysica Acta, 1509(2000)496–504.

32. Vasilyev A., Khater K., and Rakowski R.F., Effect of extracellular pH on presteady-state and steady-state current mediated by the Na/K pump, J. Membrane Biology, 198:65–76 (2004).

